# A Critical Window of Maternal Temperature Effects on Weedy Rice Seed Dormancy

**DOI:** 10.64898/2025.12.12.693925

**Authors:** Gabriela Auge, Rio Nishikata, Toshiyuki Imaizumi

**Affiliations:** Japan Society for Promotion of Science International Invitational Fellow; Consejo Nacional de Investigaciones Científicas y Tecnológicas (CONICET); Instituto de Agrobiotecnología y Biología Molecular (IABIMO), Instituto Nacional de Tecnología Agropecuaria (INTA) - CONICET. Hurlingham (CP 1686), Argentina; Institute for Plant Protection, National Agriculture and Food Research Organization (NARO), Tsukuba, 305-8604, Japan; Kyushu Okinawa Agricultural Research Center, NARO, Chikugo, 833-0041, Japan

**Keywords:** dormancy, environment, maternal effects, temperature, *Oryza sativa*, weedy rice

## Abstract

Seed dormancy is a critical trait for the survival and adaptation of agricultural weeds. The maternal temperature after flowering influences the post-dispersal response of progeny seeds, which adds layers of complexity to weed control management programs. Understanding when the maternal environment exerts its effect on seed dormancy provides valuable insights into plant fitness and weed management. In this study, we investigated weedy rice progeny seed responses to determine the sensitivity window to the maternal temperature effects in controlled and field environments. We observed that the occurrence of hot temperatures during the late reproductive stage affected weedy rice dormancy. For this effect to influence progeny seed germination, mother plants had to experience two weeks of hot treatment, especially on weeks 4 and 5 after flowering, resulting in a delay on dormancy release after seed after-ripening. Early and late transplanting in the field, which exposed mother plants to different environments after flowering, showed delayed release from dormancy consistent with the sensitivity window and responses in controlled conditions. Altogether, our results indicate that the timing and duration of the maternal environment fluctuation may serve as an input to better predict weedy rice seed responses and develop targeted and more precise control programs.

**Highlight:** Two weeks of maternal heat in the late reproductive stage are critical for delaying the release of dormancy in weedy rice seeds, which can affect plant fitness and control strategies.

## Introduction

Seed dormancy is a critical adaptive trait in the plant life cycle as it ensures germination occurs under optimal conditions for subsequent plant growth (Donohue *et al*., 2010). Seed dormancy, by regulating germination timing, is therefore essential for understanding local adaptation of plants (Postma and Ågren, 2016; Footitt *et al*., 2021; D’Aguillo and Donohue, 2023). Variation in dormancy levels is regulated by epi/genetic and environmental factors during seed maturation and after dispersal, affecting either its establishment, maintenance or release (Finch-Savage and Leubner-Metzger, 2006; Sajeev *et al*., 2024).

Seed dormancy is also regulated by the maternal environment. While developing and maturing in the mother plant, the environment exerts a strong influence on the post-dispersal response of seeds. These maternal effects can either induce or inhibit germination through the regulation of seed dormancy levels (Iwasaki *et al*., 2022). The temperature during seed development and maturation (hereafter referred to as the maternal temperature) can be considered one of the main environmental signals that control post-dispersal dormancy levels. Specifically, low maternal temperatures increase dormancy in *Avena fatua* (Peters, 1982), wheat (Reddy *et al*., 1985), *Arabidopsis thaliana* (Burghardt *et al*., 2016) and *Polygonum aviculare* (Fernández Farnocchia *et al*., 2019), whereas hot maternal temperatures increase seed dormancy in sunflower (Bodrone *et al*., 2017), cultivated rice (*Oryza sativa* L. (Ikehashi, 1972)) and weedy rice (*Oryza* spp., (Imaizumi *et al*., 2022)). Interestingly, a shift of merely 1°C in maternal temperature dramatically altered its effects on dormancy in *A. thaliana* (Springthorpe and Penfield, 2015; Huang *et al*., 2018) and weedy rice (Imaizumi *et al*., 2025). Recent studies of the maternal temperature effect revealed that it regulates seed dormancy and seedling emergence in weedy rice, as higher maternal temperature delays seedling emergence during rice cultivation (Imaizumi *et al*., 2022, 2025). The evidence shows that the experience of higher maternal temperatures can result in the potential spreading of seedling emergence times, increasing the complexity for weedy rice control in the field. Weedy rice, a conspecific to cultivated rice that has evolved through multiple independent de-domestication events (Qiu *et al*., 2020), poses a threat to rice cultivation globally (Ziska *et al*., 2015). Weedy rice populations worldwide show deeper seed dormancy than cultivated rice (Xia *et al*., 2011; Karn *et al*., 2020; Pipatpongpinyo *et al*., 2020; Imaizumi *et al*., 2022; Fukuda *et al*., 2023). Higher dormancy levels of weedy rice seeds correlate positively with seedbank longevity and negatively with seedling emergence (Pipatpongpinyo *et al*., 2020; Imaizumi *et al*., 2025), making weed occurrence in rice fields a long-term problem. For agricultural weeds, a deep seed dormancy level enhances the persistence of weed seeds in the soil seed banks (Benech-Arnold *et al*., 2000). After dispersal, dormant weed seeds replenish the soil bank, preventing germination right after dispersal in potentially unfavourable conditions. In consequence, seed dormancy is key for weed survival and adaptation to agricultural practices (Batlla *et al*., 2020).

Determining the sensitivity window of maternal temperature effects on seed dormancy is important for more accurately predict seed dormancy and seedling emergence. However, knowledge of this sensitivity window is still limited and has been mostly studied for its effects on agricultural traits in crops rather on seed dormancy. In rice, hot temperatures at the early or mid-stages of grain filling has a greater influence on grain quality than when occurring at a later stage (Tashiro and Wardlaw, 1991; Morita *et al*., 2005). Additionally, in wheat, high night temperatures during the late grain filling stage lead to reduced grain yield, grain weight, and grain weight per spike when compared to an earlier grain developmental stage (booting) (Mamrutha *et al*., 2020). In bread wheat and malting barley, raising the minimum temperature during the night both early post-flowering or from 10 days after anthesis to harvest negatively affected grain number and, consequently, grain yield (García *et al*., 2015, 2016). Raising night temperatures also influence wheat grain quality, reducing bread-making properties such as dough development and stability times, and softening degree (Li *et al*., 2024).

Despite documented effects of the maternal environment on weedy rice, when this effect is established during the reproductive stage or how long mother plants must be exposed to the cue to affect their progeny’s seed responses remain unknown. This study addresses these gaps by examining the timing of the establishment and the required duration of the cue for influencing progeny weedy rice seed responses. Our research aimed to answer the following questions: What is the sensitivity window for the maternal effect after flowering? Do the timing and length of cue exposure matter for modulating progeny responses? And finally, are the timing and duration of the maternal temperature effect relevant for weedy rice seeds developing and maturing in field conditions?

## Materials and methods

### Plant materials

In this study, we used previously described Japanese weedy rice lines (Imaizumi *et al*., 2021). These were used to test for the effect of the timing and duration of a hot treatment during the reproductive stage of mother plants. We run two growth chamber experiments, for which we chose the line ‘JP_1179’ (TEJ-derived straw hull 1; SH1_TEJ) due to its deep seed dormancy level, early flowering phenotype and short plant height that make the strain suitable for growing in a growth chamber. For the field experiment, we selected 7 weedy rice lines (3 black hull, BH; 2 SH1_TEJ; and 2 TEJ-derived straw hull 2, SH2_TEJ), namely JP_1165, JP_1177 and JP1183 for BH; JP_1179 and JP_1181 for SH1_TEJ; JP_1166 and JP_1178 for SH2_TEJ; and one cultivated rice variety (Koshihikari). The seeds for the experiments were sourced from plants grown in a paddy field in the summers of 2019 and 2021 in the Tsukuba-Kannondai test field at the National Agriculture and Food Research Organization (NARO, Japan) (36.0°N, 140.1°E).

For the first growth chamber experiment, seeds were sown in 55 mm Petri dishes filled with 5 ml of distilled water and incubated at 28°C for 2-3 days. Germinated weedy rice seeds were sown in seedling trays with rice cultivation soil (Bonsoru No. 2, Sumitomo Chemical, Japan) and placed in a greenhouse or growth chamber until transplanting in pots (∼4 weeks). One individual seedling per pot was transferred to 1/5000a Wagner pots (height 20 cm; diameter 16 cm) filled with paddy field soil fertilized with an N:P:K = 14:14:14% fertilizer at a rate of 712 mg per pot before planting. Pots were randomly arranged in growth chambers (KG-206, Koito Electric Industries Ltd., Japan) for treatments. The control growth conditions for the first experiment were set at a 28/22°C temperature cycle with a photoperiod of 12h light / 12h darkness. Plants were checked for heading and flowering on a regular basis. After flowering, the plants were separated in four contrasting temperature treatments until harvest: 1) control at 28/22°C (CC), 2) early hot treatment at 32/26°C (weeks 1, 2 and 3 after flowering, HC), 3) late hot treatment (weeks 4, 5, and 6 after flowering, CH), and 4) hot treatment control from flowering until harvest (HH) (Figure 1). The seeds were harvested as described above and kept separately in individual paper envelopes by mother plant. Six individual mother plants were grown per treatment (biological replicates). We repeated this experiment twice with similar results.

**Figure 1.**
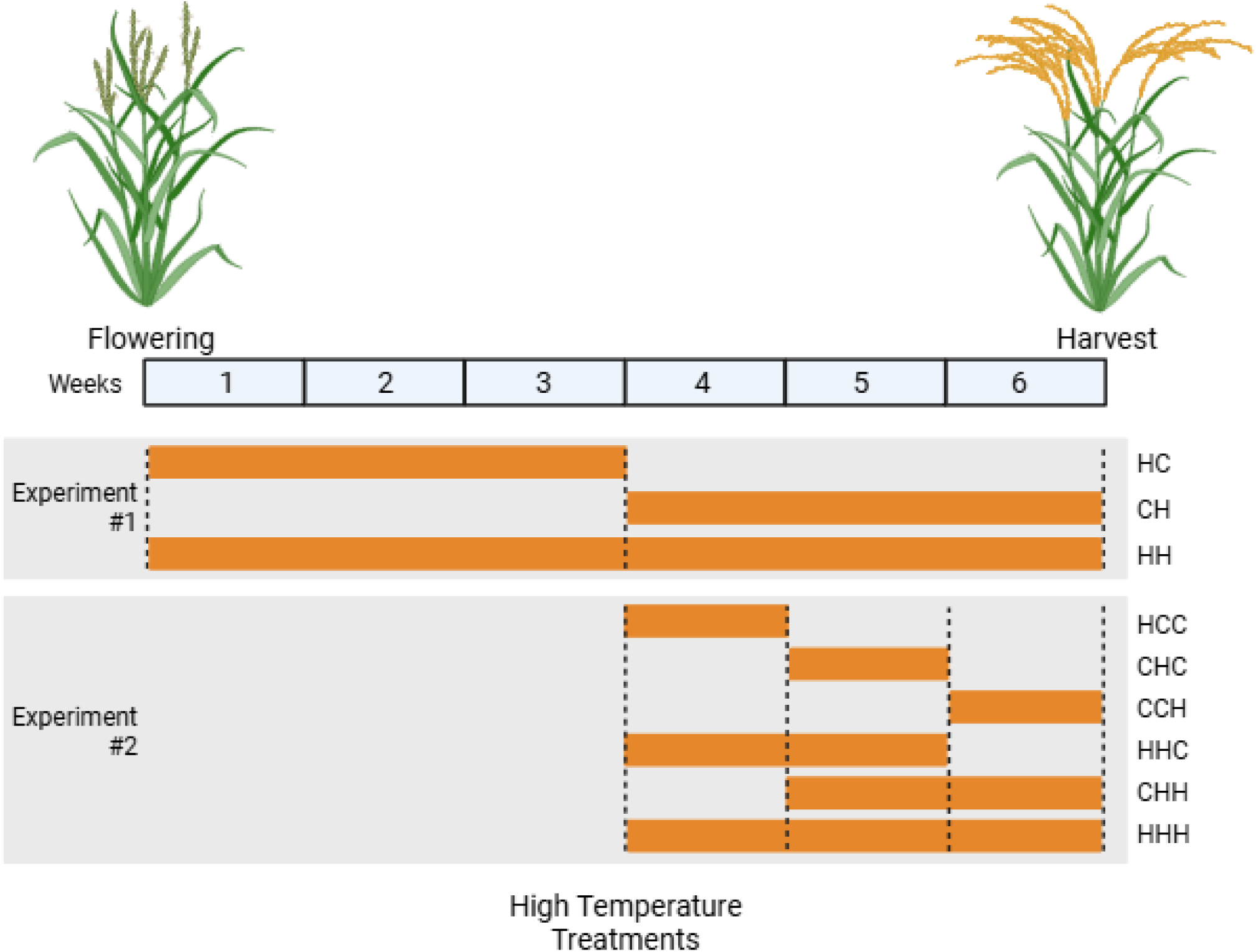
Scheme of maternal temperature treatments in controlled conditions. Orange bars indicate the occurrence of the hot temperature treatments (32/26°C day/night) during the seed development and maturation period. Gray blocks show the treatments for the first (up) and second (bottom) experiments in the growth chambers. Column to the right of the gray panels indicate the key labels to the treatments.

For the second growth chamber experiment, the plants were grown in control conditions until 3-weeks after flowering as described above. From week 4 after flowering and on, the plants were separated into the following treatments: 1) control at 28/22°C (CCC), 2) 1-week of hot treatment (32/26°C) on week 4 after flowering (HCC), 3) 1-week of hot treatment on week 5 after flowering (CHC), 4) 1-week of hot treatment on week 6 after flowering (CCH), 5) 2-weeks of hot treatment on weeks 4 and 5 after flowering (HHC), 6) 2-weeks of hot treatment on weeks 5 and 6 after flowering (CHH), and 7) 3-weeks of hot treatment control starting 3-weeks after flowering (HHH) (Figure 1). The seeds were harvested as described above and kept separately in individual paper envelopes by mother plant. Six individual mother plants were grown per treatment (biological replicates). Weeks after flowering were used as an agronomic proxy for seed developmental stage across treatments. However, because hot temperatures can accelerate development, week-based treatment windows may not perfectly align with physiological stages. Nonetheless, under our week-based timing, the temperature treatments influenced the response in JP_1179, following previous results showing the expression of maternal temperature effects (Imaizumi *et al*., 2022).

For the field experiment, seeds of the selected weedy rice strains were sown in seedling trays with commercial rice cultivation soil (Bonsoru No. 2, Sumitomo Chemical, Japan), and the trays were then placed in a greenhouse for 2-4 weeks until transplanting. Seedlings were transplanted into a paddy field on two dates: 1) May 11th 2022, and 2) June 22nd 2022. The experiments were conducted in the Tsukuba-Kannondai experimental field at the National Agriculture and Food Research Organization (NARO) (36.0°N, 140.1°E). A total of 356 kg/ha of an N:P:K = 14:14:14% was basally applied before transplanting in the field plot, and the plants were spaced at one plant per hill with 15 cm × 30 cm spacing. The plants were harvested approximately six weeks after flowering, when the seeds achieved physiological maturity. After harvest, the seeds were kept separately in individual paper envelopes by mother plant. Eight individual mother plants were grown per transplant date and line (biological replicates). The maternal temperature was calculated from air temperature data obtained from the Weather Data Acquisition System of the Institute for Agro-Environmental Sciences, NARO (https://www.naro.affrc.go.jp/org/niaes/aws/).

### Germination assays

In each experiment, fresh seeds (3 days after harvest) or seeds after-ripened for 5, 10 and 20 weeks were used for the germination assays. To facilitate detection of germination responses among treatments, we used two incubation conditions: darkness at 15°C or 30°C, accounting for suboptimal and optimal germination conditions, respectively (Fukuda *et al*., 2023). Each replicate was comprised of 20 seeds from individual mother plants sown in 55 mm Petri dishes filled with 5 ml of distilled water. The plate positions in the germination chambers were randomized within each incubation treatment after each germination census to minimize position effects.

Germination was scored at 4, 7, 10 and 14 days after sowing. A seed was considered germinated when the protrusion of the radicle from the seed coat was visible. Germinated seeds were removed from the Petri dishes at each census. Germination reached a clear plateau after 14 days of incubation. Seeds were considered viable when firm after pressed with a pestle. We recorded mouldy or mushy seeds as not viable and removed them from the plates during the incubation period.

### Statistical analysis

All analyses were performed using R ver. 4.4.1 (R Development Core Team, 2024). To evaluate the maternal temperature effects on progeny seed germination, the final germination proportions were analyzed with generalized linear models with a quasibinomial distribution and a logit link using the base package. For the growth chamber experiments, we first run a full model including final germination rate as the response variable and maternal treatment (Treatment), after-ripening time (AR time) and incubation temperature (Inc Temp) as fixed factors. For the field experiment, the full model included transplanting date (Transplant), weedy rice group (Type), after-ripening time (AR time) and incubation temperature (Inc Temp) as explanatory fixed variables. After-ripening time was considered a continuous variable, while maternal treatment (Treatment and Transplant), weedy rice group (Type) and incubation temperature (Inc Temp) as categorical. In all cases, we first analyzed the general variable effects and their interactions. Analysis of deviance based on F-values was performed using the function “Anova” in the “car” (v3.0.10) package (Fox and Weisberg, 2010). We also run submodels by incubation temperature to interpret germination responses in the contrasting sub- and optimal germination temperatures. We visualized germination after different periods of dry storage (after-ripening time) and estimated the time to reach 50% germination (G50) and the odds ratio using the estimated values and 95% confidence intervals (CIs) from the model. We used the G50 values to estimate the level of dormancy.

## Results

### Sensitivity window of maternal temperature effects

Hot temperatures during the reproductive stage of the mother plants delay seed dormancy release (Imaizumi *et al*., 2022). We first tested whether this effect requires the persistent experience of hot conditions throughout the whole period of seed development and maturation. We compared germination responses of progeny seeds harvested from plants either grown in control conditions for the first 3 weeks after flowering and then moved to the hot treatment until harvest (28/22°C -> 32/26°C:CH) or viceversa (32/26°C -> 28/22°C: HC) with those from plants grown in control conditions (CC) or the hot treatment from flowering to harvest (HH). Fresh seeds from all treatments showed a high level of dormancy (Figure 2). With after-ripening progressing, seeds from mother plants grown in control conditions (CC) lost dormancy faster than those grown at hot temperatures during the whole reproductive stage (HH) at both incubation temperatures (significant Treatment x AR and Treatment x Inc Temp, Supp. Table 1A), confirming our treatments were yielding the desirable results (Figure 2; (Imaizumi *et al*., 2022)). Seeds from the CC treatment mother plants needed 15.7 (95% CI = 14.3-17.3) and 7.4 weeks (6.2-8.5) of after-ripening time to reach 50% germination (G50) compared with >20 (low CI > 20) and 11.6 weeks (8.1-16.6) for the HH treatment seeds when incubated at 15 and 30°C, respectively (Table 1). Germination odds ratios for the HH compared to CC treatment were 0.87 (0.82–0.92) and 0.86 (0.82–0.90) for seeds incubated at 15°C and 30°C, respectively (Supp. Table 2), supporting that seeds from the HH condition required longer time to release dormancy.

**Figure 2.**
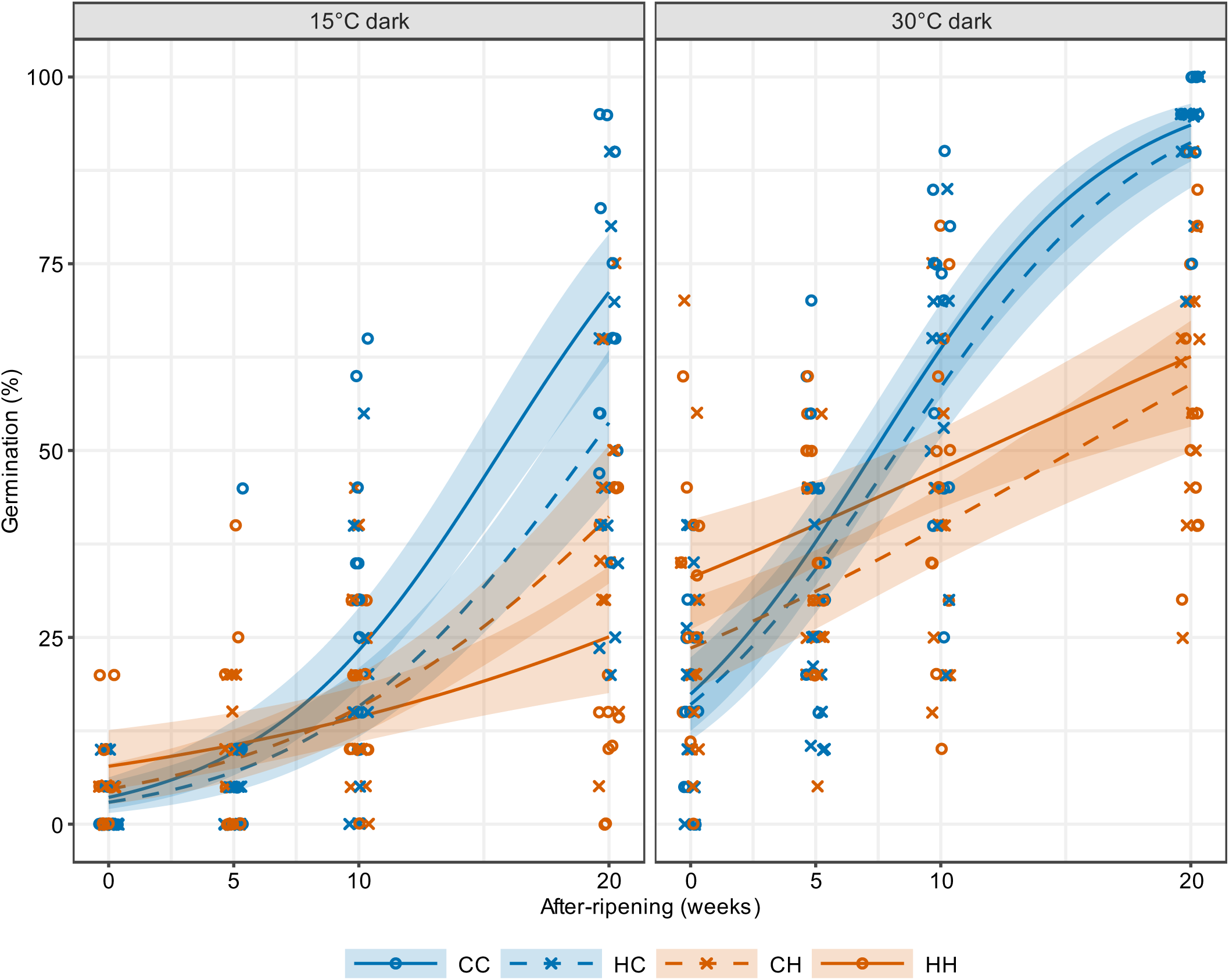
Weedy rice progeny seed germination response after maternal temperature treatments. Germination proportion along the course of after-ripening (x-axis) were estimated using a quasibinomial generalized linear model. Each line represents the maximum likelihood estimates for each maternal treatment (solid blue line: control, CC; dashed blue line: early hot, HC; solid orange line: hot control, HH; and dashed orange line: late hot, CH), and shaded areas represent the 95% confidence intervals for each estimate. Symbols indicate final germination proportion for each replicate at each after-harvest time point.

**Table 1.** After-ripening weeks required to reach 50% germination (G50) for the first experiment in controlled conditions. The time to reach G50 and the confidence intervals were estimated for each treatment (CC: control maternal temperature; CH: later hot maternal treatment; HC: early hot maternal treatment; and HH: control hot maternal treatment) for progeny seeds incubated at 15 and 30°C.

| Temperature | Treatment | G50 | Lower CI (2.5%) | Upper CI (97.5%) |
| --- | --- | --- | --- | --- |
| 15°C | CC | 15.7 | 14.3 | 17.3 |
|  | CH | > 20 weeks | 19.8 | > 20 weeks |
|  | HC | 19.2 | 17.4 | > 20 weeks |
|  | HH | > 20 weeks | > 20 weeks | > 20 weeks |
| 30°C | CC | 7.4 | 6.2 | 8.5 |
|  | CH | 15.3 | 12.4 | > 20 weeks |
|  | HC | 8.3 | 7 | 9.6 |
|  | HH | 11.6 | 8.1 | 16.6 |

When the hot treatment was applied early in the reproductive stage (HC), the seeds displayed a germination pattern similar to that of control seeds (CC), with after-ripening time to reach G50 also similar to that of CC seeds (although slightly higher; Table 1). Interestingly, the seeds from mother plants that experienced the hot treatment late (CH) showed a similar germination response and number of after-ripening weeks to reach G50 to that of seeds from plants grown in the HH treatment (Figure 2, Table 1). The germination odds ratio compared to those of the control condition (CC) were lower in the CH treatment and similar to the HH treatment (Supp. Table 2). However, odds were not different for the early hot treatment HC (0.97, CI = 0.92–1.03, and 0.99, CI = 0.93–1.05, at 15°C and 30°C incubation temperatures, respectively).

### The maternal effect depends on the timing and duration of the stimulus

Average mean daily temperatures experienced by plants during the reproductive stage for the last 40 years (estimated by transplanting and flowering times for the lines used in this study) exhibit a wider range of variance when assessed weekly as opposed to every two or three weeks (Supp. Figure 1), indicating larger variation in shorter times may result in higher yearly variation in dormancy levels in progeny seeds. To assess whether the timing and duration of the maternal environment effect influences the response of the progeny seeds, we subjected the plants to the hot treatment for 1, 2 or 3-weeks during the sensitivity window we have established in the previous experiment (weeks 4, 5 and 6 after flowering) (Figure 2). The maternal environment had a significant effect on progeny germination that depended on the time after harvest (significant Treatment x AR time; Supp. Table 1B). Progeny seeds from mothers that experienced only 1 week of hot treatment (HCC, CHC, and CCH) showed a similar response to those of the CCC control treatment, regardless of when that treatment was applied (Figure 3; Supp. Figure 2). Seeds of CCC control plants and those from plants experiencing 1 week of the hot treatment (HCC, CHC, and CCH) that were incubated at 15°C required 14-16 weeks of after-ripening to reach G50, while those incubated at 30°C required 8-10 weeks (Table 2). The hot maternal environment was required to be experienced for 2-weeks (HHC and CHH) to significantly delay dormancy release in the progeny, showing a similar effect to that of 3-weeks of hot temperature within the sensitivity window (HHH treatment) (Figure 3). Two-weeks of hot temperature delayed the number of after-ripening weeks to reach G50 to more than 20 weeks for seeds incubated at 15°C and 11 weeks for those incubated at 30°C, similar to the response of seeds from the late hot treatment control (HHH; > 20 and 12.2 weeks for seeds incubated at 15 and 30°C, respectively) (Table 2). Although the effect is similar at both incubation temperatures, germination patterns of seeds incubated at 30°C (optimal incubation temperature) showed slightly larger effects of the maternal temperature on dormancy release (Supp. Table 2): the germination odds ratio for the hot treatment experienced only in week 6 (CCH) was 0.89 (CI = 0.79–0.97), which was as low as the odds ratio of progeny seeds from plants exposed to hot temperatures for two (HHC and CHH) or three weeks (HHH) (0.86-0.9).

**Figure 3.**
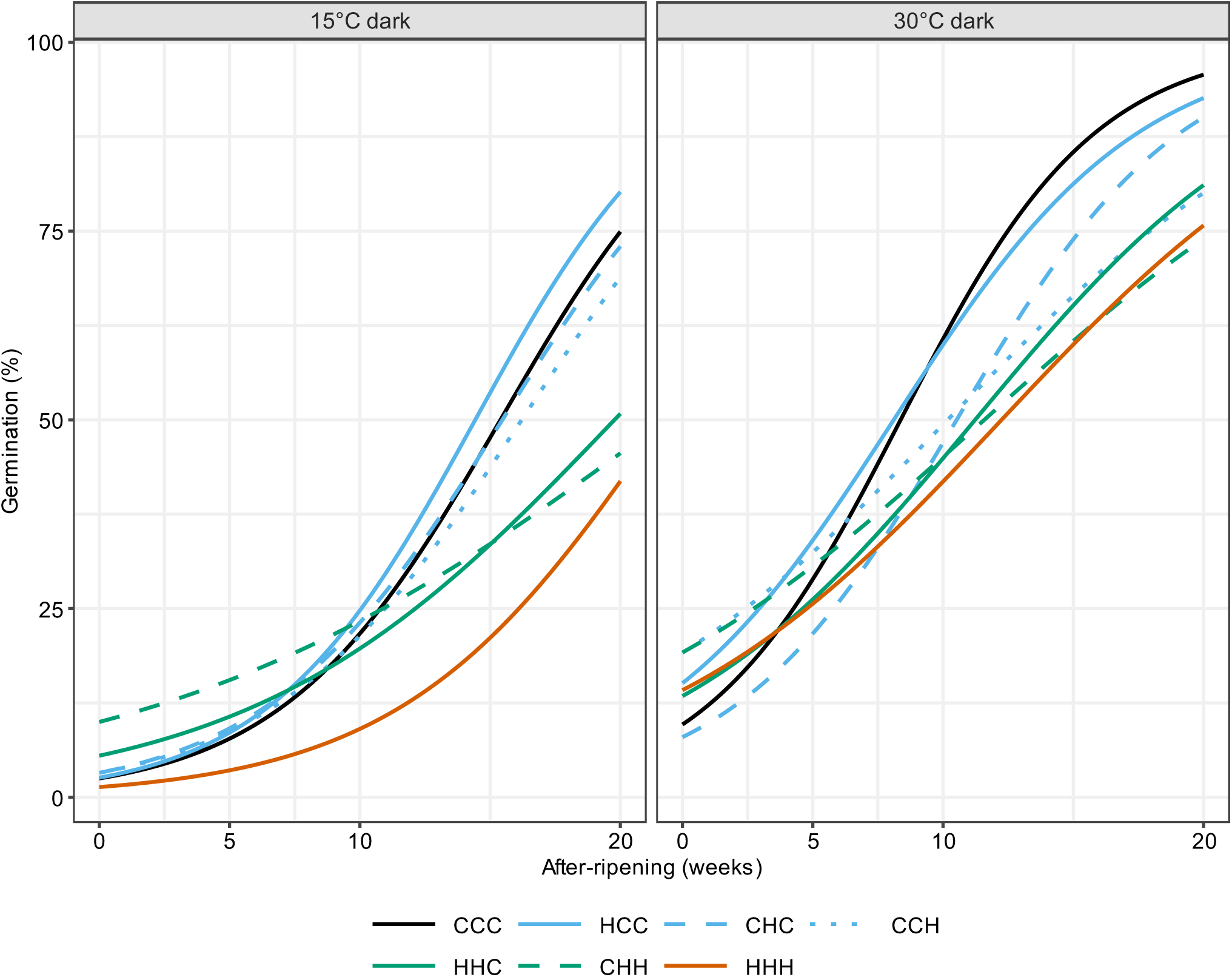
Weedy rice progeny seed germination response after maternal temperature treatments to test for timing and duration of the stimulus. Germination rates during the course of after-ripening (x-axis) were estimated using a quasibinomial generalized linear model. Each line represents the maximum likelihood estimates for each maternal treatment (solid black line: control treatment, CCC; solid blue line: 1 week of hot treatment on week 4 after flowering, HCC; dashed blue line: 1 week of hot treatment on week 5 after flowering, CHC; dotted blue line: 1 week of hot treatment on week 6 after flowering, CCH; solid green line: 2 weeks of hot treatment on weeks 4 and 5 after flowering, HHC; dashed green line: 2 weeks of hot treatment on weeks 5 and 6 after flowering, CHH; and solid orange line: 3 weeks of hot treatment on weeks 4, 5 and 6, HHH). See also Supplementary Figure 2.

**Table 2.** After-ripening weeks required to reach 50% germination (G50) for the second experiment in controlled conditions. The time to reach G50 and the confidence intervals were estimated for each treatment (CCC: control treatment; HCC: 1 week of hot treatment on week 4 after flowering; CHC: 1 week of hot treatment on week 5 after flowering; CCH: 1 week of hot treatment on week 6 after flowering; HHC: 2 weeks of hot treatment on weeks 4 and 5 after flowering; CHH: 2 weeks of hot treatment on weeks 5 and 6 after flowering; and HHH: 3 weeks of hot treatment on weeks 4, 5 and 6) for progeny seeds incubated at 15 and 30°C.

| Temperature | Treatment | G50 | Lower CI (2.5%) | Upper CI (97.5%) |
| --- | --- | --- | --- | --- |
| 15°C | CCC | 15.4 | 13.5 | 17.8 |
|  | HCC | 14.4 | 12.6 | 16.6 |
|  | CHC | 15.5 | 13.5 | 18.1 |
|  | CCH | 16.2 | 14.1 | 19.0 |
|  | HHC | 19.8 | 16.6 | > 20 weeks |
|  | CHH | > 20 weeks | 17.1 | > 20 weeks |
|  | HHH | > 20 weeks | 18.9 | > 20 weeks |
| 30°C | CCC | 8.4 | 6.9 | 10.1 |
|  | HCC | 8.1 | 6.3 | 10.0 |
|  | CHC | 10.5 | 8.8 | 12.5 |
|  | CCH | 10.2 | 7.7 | 13.1 |
|  | HHC | 11.2 | 9.1 | 13.9 |
|  | CHH | 11.6 | 8.9 | 15.2 |
|  | HHH | 12.2 | 9.9 | 15.4 |

### Differences in progeny germination response in the field

To evaluate whether the maternal effects as described above can be detected in the field, we run an experiment in which we transplanted seedlings in two different dates (mid-May and late-June) resulting in two different maternal temperature environments after flowering. We observed that seeds from plants transplanted in mid-May showed a delayed dormancy release compared with those from mother plants transplanted in late-June, and overlaps with higher daily mean temperatures during the sensitivity window (Figure 4). Temperatures experienced by plants differed between the weedy rice groups due to their flowering dates (Supp. Table 3), but plants of the SH1 group transplanted in May were the only ones experiencing daily mean temperatures above 27°C on weeks 4 and 5, which correlates with the sensitivity period and response detected in controlled conditions (Figure 4B). This temperature has been reported as the threshold for maternal effects in seedling emergence in the field (Imaizumi *et al*., 2025). For this group, the number of after-ripening weeks needed to reach G50 were 6.2 and 5.4 for seeds incubated at 30°C of plants transplanted in May and June, respectively (Table 3). On the other hand, seeds of SH2 plants experiencing a temperature above 27°C only on week 4 showed a reduced maternal effect on germination (no difference in after-ripening weeks to reach G50 between transplanting dates; Table 3). However, for seeds of the BH group, only one week with daily mean temperatures of ∼27°C was enough to induce a maternal effect on germination (5.8 and 3.8 after-ripening weeks to reach G50 when seeds were incubated at 30°C, and 19 and 14.7 weeks for those incubated at 15°C, respectively for May and June transplanted plants; Table 3), suggesting a higher sensitivity to the environment as we have previously shown (Imaizumi *et al*., 2022) (Figure 4).

**Figure 4.**
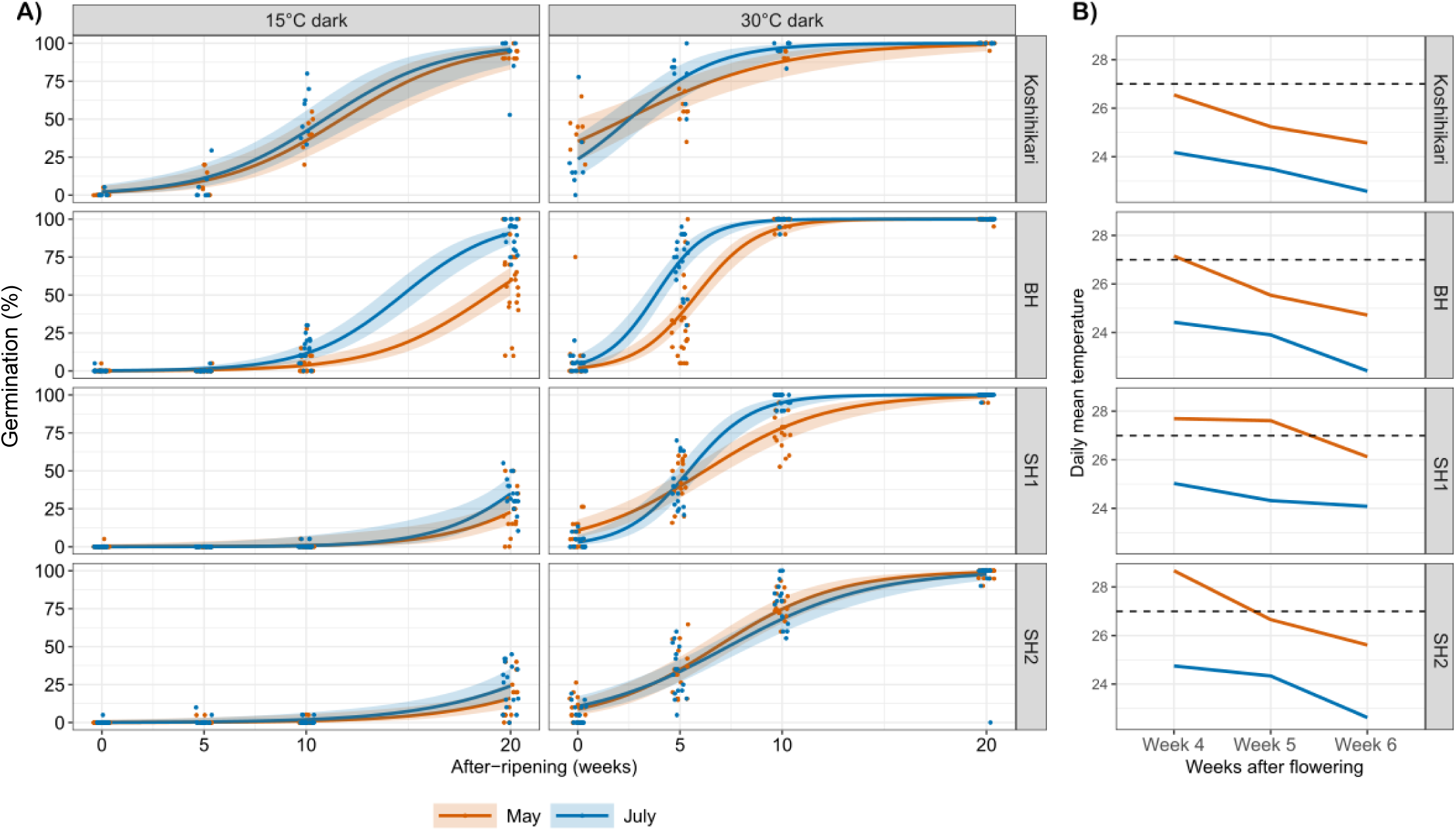
Weedy rice progeny seed germination response after maternal temperature treatments due to different transplanting dates in the field. A) Germination rates along the course of after-ripening (x-axis) were estimated using a quasibinomial generalized linear model for seeds of the BH, SH1 and SH2 weedy rice types, and the Koshihikari cultivar. Each line represents the maximum likelihood estimates for each maternal treatment (blue line: seeds from mother plants transplanted in mid-May, and orange: seeds from mother plants transplanted in late-June), and shaded areas represent the 95% confidence intervals for each estimate. Symbols indicate final germination proportions for each replicate at each after-harvest time point. B) Calculated daily mean temperature experienced on weeks 4, 5 and 6 after flowering for plants transplanted in mid-May (orange) and late-June (blue) for each weedy rice group. The black dashed line indicates the temperature threshold for the establishment of maternal effects (Imaizumi *et al*., 2025)

**Table 3.** After-ripening weeks required to reach 50% germination (G50) for the field experiment. The time to reach G50 and the confidence intervals were estimated for each transplanting date (May and June) and weedy rice group (Koshihikari; black hull, BH; straw hull 1, SH1; and straw hull 2, SH2; see also Materials and Methods) for progeny seeds incubated at 15 and 30°C.

| Temperature | Group | Transplant Date | G50 | Lower CI (2.5%) | Upper CI (97.5%) |
| --- | --- | --- | --- | --- | --- |
| 15°C | Koshihikari | May | 11.7 | 10.1 | 13.7 |
|  | Koshihikari | June | 10.9 | 9.4 | 12.9 |
|  | BH | May | 19 | 18 | >20 |
|  | BH | June | 14.7 | 13.7 | 15.8 |
|  | SH1 | May | >20 | >20 | >20 |
|  | SH1 | June | >20 | >20 | >20 |
|  | SH2 | May | >20 | >20 | >20 |
|  | SH2 | June | >20 | >20 | >20 |
| 30°C | Koshihikari | May | 2.3 | 0 | 4 |
|  | Koshihikari | June | 2.5 | 1.2 | 3.7 |
|  | BH | May | 5.8 | 5.3 | 6.4 |
|  | BH | June | 3.8 | 3.2 | 4.3 |
|  | SH1 | May | 6.2 | 5.2 | 7.2 |
|  | SH1 | June | 5.4 | 4.8 | 6.1 |
|  | SH2 | May | 6.8 | 5.8 | 7.9 |
|  | SH2 | June | 7.4 | 6.3 | 8.5 |

## Discussion

The maternal temperature is a relevant cue for post-dispersal seed germination responses for many species (Iwasaki *et al*., 2022). In agricultural weeds, the maternal effect has direct consequences on seed dormancy levels affecting soil seed bank dynamics and seedling emergence (Imaizumi *et al*., 2025), and contributes to their adaptation to agricultural practices (Darmency *et al*., 2017; Batlla *et al*., 2020). In this study, we explored the effect of the maternal temperature environment after flowering on the germination response of the progeny to detect the sensitivity window (timing) and duration of the cue needed for those effects to be established in weedy rice. We found that the experience of hot temperatures for two weeks during the late reproductive stage is enough to influence the response of the progeny. Visualizing yearly temperature variation at a more detailed weekly time scale rather than monthly within the sensitivity window, uncovers greater fluctuations, providing valuable information on environmental patterns that can directly influence and help forecast weed emergence in future crop cycles.

We have previously shown that hot temperatures during seed maturation in the mother plant reduces the ability to release a dormant state in weedy rice (Imaizumi *et al*., 2022). In this study, we established the sensitivity window of the maternal temperature effects as the late reproductive stage period (weeks 4, 5 and 6 after flowering). An increase in temperature in this period led to a delay in dormancy release across the course of after-ripening under dry seed storage at room temperature, showing a low proportion of weedy rice seed germination even in optimal seed incubation conditions. The critical window for the establishment of the maternal temperature effect on seed dormancy was previously shown in cultivated rice: seeds developed under hot temperatures in the later stage of maturation showed a high degree of primary dormancy (Ikehashi, 1967, 1972). The maternal temperature effect on primary dormancy was weak in our study probably due to the deeper seed dormancy levels of weedy rice than cultivated rice seeds (Imaizumi *et al*. 2022), but the effect on dormancy release was strong. However, both results go in line and show that the later reproductive stage is key to perceive the environmental cue regulating progeny responses.

Dormancy levels in seeds are regulated mainly by the hormone abscisic acid (ABA) (Finch-Savage and Leubner-Metzger, 2006); thus, the effect of the maternal temperature in the sensitivity window is likely mediated by changes in ABA profiles. Hot temperature during grain filling disrupts the homeostasis of the hormone profile in developing grains and the flag leaf of the ‘Xiangzaoxian No45 (XZX 45)’ rice cultivar (Shao *et al*., 2021). ABA accumulation in the late grain filling stage (14 and 21 days after flowering) is increased, leading to a reduced mobilization of resources from the flag leaf, reduced number of grains per year and increased grain chalkiness (Shao *et al*., 2021). Hot temperature during grain filling is also related to delayed seed germination in the ‘Nipponbare’ rice cultivar, which is associated with increased expression of ABA synthesis genes, and down-regulation of ABA catabolism (*OsABA8ox3*) and amylase genes during seed imbibition in response to the maternal environment (Suriyasak *et al*., 2020). Regulation of the ABA content in seeds by the maternal temperature during the late maturation stage was also confirmed in winter annuals, such as *Brassica oleracea*. In *B. oleracea*, cold temperatures led to ABA accumulation at maturity in seeds due to slower ABA catabolism relative to synthesis, especially in the endosperm (Chen *et al*., 2021). Thus, at lower temperatures, seeds do not break primary dormancy prior to shedding. The effect of temperature is associated with lower expression of ABA metabolism genes during seed maturation in both cultivated rice and *B. oleracea* (Suriyasak *et al*., 2020; Chen *et al*., 2021). Whether overlapping regulation of ABA accumulation and maternal temperature effects in weedy rice results in variation in dormancy release in the conditions tested in this study, warrants further investigation.

We also confirmed that the sensitivity window for the maternal temperature effect is relevant for the progeny of field-grown plants. Previous work has shown that the maternal temperature effects are limited below a threshold of 27-28°C, while they were consistent among different weedy rice groups (Imaizumi *et al*., 2025). In this study, we observed that seeds matured under early summer showed a delayed dormancy release compared with those matured under late summer. Even though we have not accounted for any other environmental factor effects, the response of the seeds shows a strong pattern of correlation with higher daily mean temperatures (above 27°C) during the sensitivity window due to increased air temperature in early summer. Interestingly, seeds of the BH weedy rice group showed a delayed dormancy release than the other groups when exposed to hot temperatures for only one week, suggesting a higher sensitivity to the environment, as we have previously shown (Imaizumi *et al*., 2022). This goes in line with the observation in the second experiment in controlled conditions where a one-week exposure in week 6 produced a modest effect when seeds were incubated at 30°C. Altogether, the results indicate a strong potential of the maternal temperature to direct progeny responses.

A delayed dormancy release makes weedy rice more prone to emerge later in the year and successive cultivation cycles. Higher dormancy due to the influence of maternal temperatures have been shown to delay seedling emergence even in favorable conditions (springtime; Imaizumi *et al*., 2025). Temporal autocorrelation of selective environmental cues, such as those projected for climate change, favors the expression of adaptive traits across generations (Auge *et al*., 2023; Kanomanyanga *et al*., 2025). Germination timing strongly influences plant fitness (Donohue *et al*., 2005; Akiyama and Ågren, 2014; Postma and Ågren, 2016). Thus, identifying the sensitivity window for maternal temperature effects on seed dormancy is essential for understanding plant fitness and predicting its response to climate change. Our results show that changing maternal temperature even for a short period leads to a strong effect on variation of dormancy levels and release in weedy rice, and that this interacts with genetic variation in the field, which emerges as an adaptive mechanism to cope with future changing climates and agricultural practices.

One key goal in weed management is to predict weed seed responses efficiently in order to implement more effective control strategies. The results of our study and the previous evidence show that the experience of higher maternal temperatures can result in the potential spreading of seedling emergence times. Thus, understanding the influence of environmental cues that affect weed seed dormancy is of utmost importance. The relevance of our results extends beyond agricultural weed control, since determining the maternal temperature sensitivity window for seed dormancy is crucial for crops because weak dormancy can raise the risk of preharvest sprouting while strong dormancy may impede uniform germination and successful stand establishment after sowing.

## Supplementary Data

**Supplementary Table 1**. Effects of the maternal environment (Treatment), time after harvest (AR time) and incubation temperature (Inc Temp), and their interactions, on the response of weedy rice progeny seeds in the first and second experiments in controlled conditions.

**Supplementary Table 2.** Effect size of the maternal temperature environment in both experiments done in controlled conditions.

**Supplementary Table 3.** Flowering time and date for the weedy rice plants grown in the field.

**Supplementary Figure 1**. Average temperature variation since 1980 for the experimental field in Tsukuba, Ibaraki.

**Supplementary Figure 2.** Weedy rice progeny seed germination response after maternal temperature treatments to test for timing and duration of the stimulus.

## Acknowledgements

The authors thank the staff members of the Weed Management Group, Institute for Plant Protection, NARO and those of the Department of Technical Support of NARO for their technical assistance.

## Author contributions

GA and TI: conceptualization; GA, RT and TI: methodology; GA and TI: formal analysis; GA, RT and TI: investigation; TI: resources; GA and TI: writing - original draft; GA, RT and TI: writing - review & editing; GA and TI: funding acquisition.

## Data availability

The data that support the findings of this study are available upon request from the corresponding author.

## References

Akiyama R, Ågren J. 2014. Conflicting selection on the timing of germination in a natural population of *Arabidopsis thaliana*. Journal of Evolutionary Biology 27, 193–199.

Auge G, Hankofer V, Groth M, Antoniou-Kourounioti R, Ratikainen I, Lampei C. 2023. Plant environmental memory: implications, mechanisms and opportunities for plant scientists and beyond. AoB PLANTS 15, plad032.

Batlla D, Ghersa CM, Benech-Arnold RL. 2020. Dormancy, a critical trait for weed success in crop production systems. Pest Management Science 76, 1189–1194.

Benech-Arnold RL, Sánchez RA, Forcella F, Kruk BC, Ghersa CM. 2000. Environmental control of dormancy in weed seed banks in soil. Field Crops Research 67, 105–122.

Bodrone MP, Rodríguez MV, Arisnabarreta S, Batlla D. 2017. Maternal environment and dormancy in sunflower: The effect of temperature during fruit development. European Journal of Agronomy 82, Part A, 93–103.

Burghardt LT, Edwards BR, Donohue K. 2016. Multiple paths to similar germination behavior in *Arabidopsis thaliana*. New Phytologist 209, 1301–1312.

Chen C, Travis AJ, Hossain M, Islam MR, Price AH, Norton GJ. 2021. Genome-wide association mapping of sodium and potassium concentration in rice grains and shoots under alternate wetting and drying and continuously flooded irrigation. Theoretical and Applied Genetics 134, 2315–2334.

D’Aguillo M, Donohue K. 2023. Changes in phenology can alter patterns of natural selection: the joint evolution of germination time and postgermination traits. New Phytologist 238, 405–421.

Darmency H, Colbach N, Le Corre V. 2017. Relationship between weed dormancy and herbicide rotations: implications in resistance evolution. Pest Management Science 73, 1994–1999.

Donohue K, Dorn L, Griffith C, Kim E, Aguilera A, Polisetty CR, Schmitt J. 2005. The Evolutionary Ecology of Seed Germination of *Arabidopsis thaliana*: Variable Natural Selection on Germination Timing. Evolution 59, 758–770.

Donohue K, Rubio de Casas R, Burghardt L, Kovach K, Willis CG. 2010. Germination, Postgermination Adaptation, and Species Ecological Ranges. Annual Review of Ecology, Evolution, and Systematics 41, 293–319.

Fernández Farnocchia RB, Benech-Arnold RL, Batlla D. 2019. Regulation of seed dormancy by the maternal environment is instrumental for maximizing plant fitness in *Polygonum aviculare*. Journal of Experimental Botany 70, 4793–4806.

Finch-Savage WE, Leubner-Metzger G. 2006. Seed dormancy and the control of germination. New Phytologist 171, 501–523.

Footitt S, Hambidge AJ, Finch-Savage WE. 2021. Changes in phenological events in response to a global warming scenario reveal greater adaptability of winter annual compared with summer annual arabidopsis ecotypes. Annals of Botany 127, 111–122.

Fox J, Weisberg S. 2010. An R companion to applied regression. Los Angeles, USA: Sage Publications.

Fukuda M, Imaizumi T, Koarai A. 2023. Seed germination responses to temperature and water availability in weedy rice. Pest Management Science 79, 870–880.

García GA, Dreccer MF, Miralles DJ, Serrago RA. 2015. High night temperatures during grain number determination reduce wheat and barley grain yield: a field study. Global Change Biology 21, 4153–4164.

García GA, Serrago RA, Dreccer MF, Miralles DJ. 2016. Post-anthesis warm nights reduce grain weight in field-grown wheat and barley. Field Crops Research 195, 50–59.

Huang Z, Footitt S, Tang A, Finch□Savage WE. 2018. Predicted global warming scenarios impact on the mother plant to alter seed dormancy and germination behaviour in *Arabidopsis*. Plant, Cell & Environment 41, 187–197.

Ikehashi H. 1967. Studies on the enviromental fluctuation of germination habits of rice seeds and the test and selection methods for them. I. On the effect of temperature during maturation on germination of rice seeds. Japanese Journal of Breeding 17, 72–77.

Ikehashi H. 1972. Induction and test of dormancy of rice seeds by temperature condition during maturation. Japanese Journal of Breeding 22, 209–216.

Imaizumi T, Ebana K, Kawahara Y, Muto C, Kobayashi H, Koarai A, Olsen KM. 2021. Genomic divergence during feralization reveals both conserved and distinct mechanisms of parallel weediness evolution. Communications Biology 4, 1–11.

Imaizumi T, Kawahara Y, Auge G. 2022. Hybrid-derived weedy rice maintains adaptive combinations of alleles associated with seed dormancy. Molecular Ecology 31, 6556–6569.

Imaizumi T, Ohigashi K, Koarai A. 2025. Maternal Temperature Imposes a Longer-Term Effect on Seedling Emergence Than Does Genetic Variation in Seed Dormancy. Plant, Cell & Environment 48, 5304–5316.

Iwasaki M, Penfield S, Lopez-Molina L. 2022. Parental and Environmental Control of Seed Dormancy in Arabidopsis thaliana. Annual Review of Plant Biology 73, 355–378.

Kanomanyanga J, Cussans J, Moss S, Ober E, Liu C, Coutts S. 2025. Adaptation of grassweeds to spring cropping through changes in germination, flowering time and fecundity. Scientific Reports 15, 21492.

Karn E, Leon TD, Espino L, Al-Khatib K, Brim-DeForest W. 2020. Phenotypic diversity of weedy rice (*Oryza sativa* f. spontanea) biotypes found in California and implications for management. Weed Science 68, 485–495.

Li D, Xiao Y, Guo L, et al. 2024. Effect of High Nighttime Temperatures on Growth, Yield, and Quality of Two Wheat Cultivars During the Whole Growth Period. Plants 13, 3071.

Mamrutha HM, Rinki K, Venkatesh K, Gopalareddy K, Khan H, Mishra CN, Kumar S, Kumar Y, Singh G, Singh GP. 2020. Impact of high night temperature stress on different growth stages of wheat. Plant Physiology Reports 25, 707–715.

Morita S, Yonemaru JI, Takanashi JI. 2005. Grain Growth and Endosperm Cell Size Under High Night Temperatures in Rice (*Oryza sativa* L.). Annals of Botany 95, 695–701.

Peters NCB. 1982. The dormancy of wild oat seed (*Avena fatua* L.) from plants grown under various temperature and soil moisture conditions. Weed Research 22, 205–212.

Pipatpongpinyo W, Korkmaz U, Wu H, Kena A, Ye H, Feng J, Gu XY. 2020. Assembling seed dormancy genes into a system identified their effects on seedbank longevity in weedy rice. Heredity 124, 135–145.

Postma FM, Ågren J. 2016. Early life stages contribute strongly to local adaptation in *Arabidopsis thaliana*. Proceedings of the National Academy of Sciences 113, 7590–7595.

Qiu J, Jia L, Wu D, et al. 2020. Diverse genetic mechanisms underlie worldwide convergent rice feralization. Genome Biology 21, 70.

R Development Core Team. 2024. R: A language and environment for statistical computing. Vienna: R Foundation for Statistical Computing, Vienna, Austria.

Reddy LV, Metzger RJ, Ching TM. 1985. Effect of Temperature on Seed Dormancy of Wheat. Crop Science 25, 455–458.

Sajeev N, Koornneef M, Bentsink L. 2024. A commitment for life: Decades of unraveling the molecular mechanisms behind seed dormancy and germination. The Plant Cell, koad328.

Shao C, Shen L, Qiu C, et al. 2021. Characterizing the impact of high temperature during grain filling on phytohormone levels, enzyme activity and metabolic profiles of an early indica rice variety. Plant Biology 23, 806–818.

Springthorpe V, Penfield S. 2015. Flowering time and seed dormancy control use external coincidence to generate life history strategy. eLife, e05557.

Suriyasak C, Oyama Y, Ishida T, Mashiguchi K, Yamaguchi S, Hamaoka N, Iwaya-Inoue M, Ishibashi Y. 2020. Mechanism of delayed seed germination caused by high temperature during grain filling in rice (*Oryza sativa* L.). Scientific Reports 10, 17378.

Tashiro T, Wardlaw IF. 1991. The effect of high temperature on kernel dimensions and the type and occurrence of kernel damage in rice. Australian Journal of Agricultural Research 42, 485–496.

Xia HB, Xia H, Ellstrand NC, Yang C, Lu BR. 2011. Rapid evolutionary divergence and ecotypic diversification of germination behavior in weedy rice populations. New Phytologist 191, 1119–1127.

Ziska LH, Gealy DR, Burgos N, et al. 2015. Chapter Three - Weedy (Red) Rice: An Emerging Constraint to Global Rice Production. In: Sparks DL, ed. Advances in Agronomy. Academic Press, 181–228.

